# *rapunzel5* is necessary for normal hematopoietic development in zebrafish

**DOI:** 10.1101/2025.08.09.669486

**Authors:** S Thapa, W Dowell, E Harris, DL Stachura

## Abstract

The molecular mechanisms regulating the highly complex process of hematopoiesis in vertebrates is still enigmatic. This system begins with the controlled differentiation of adult hematopoietic stem and progenitor cells (HSPCs), which can proliferate and generate all types of mature blood cells. Identifying the underlying factors and mechanisms that allow HSPCs to differentiate and proliferate is an essential issue in stem cell biology and tissue homeostasis; disruptions in these processes can cause severe diseases. A transcriptomic screen of hematopoietic-supportive zebrafish stromal tissues identified *rapunzel5 (rpz5)* was likely involved in vertebrate hematopoiesis. We performed loss-of-function experiments in zebrafish embryos at the one-cell-stage with morpholinos to determine if blood cell production was affected by *rpz5*. *rpz5* knockdown resulted in reduced amounts of red blood cells, myeloid cells, and thrombocytes, and adding back exogenous *rpz5* rescued these deficiencies. Further analysis with methylcellulose assays indicated that there was also a significant reduction in HSPCs following *rpz5* reduction. Together, these findings suggest that zebrafish *rpz5* is essential for normal formation, differentiation, and proliferation of HSPCs, specifically down the erythroid and myeloid pathway. Fully understanding the roles of novel genes such as *rpz5* is essential for understanding the evolution of vertebrate hematopoiesis and for treating hematological diseases in the future.

## Introduction

Hematopoiesis is the development of blood cells that an organism needs during its whole life span. At the core of this system are hematopoietic stem cells (HSCs), which have the unique property of self-renewal and differentiation down the myeloid and lymphoid pathways. This allows HSCs to constantly replenish themselves while simultaneously creating and maintaining red blood cells and the cells of the innate and adaptive immune system. This complex process of hematopoiesis is regulated by various molecular pathways that affect hematopoietic stem and progenitor cell (HSPC) growth, differentiation, proliferation, and homeostasis. When this process becomes disordered, multiple types of blood disorders can occur, such as anemia, thrombocytopenia, leukopenia, as well as blood cancers (leukemia). While the hematopoietic system has been studied for decades, molecular pathways are still being discovered that regulate the process of blood production. These pathways help us understand not only the evolution of the hematopoietic system, but also potential mechanisms that could be exploited for therapeutic use in the future.

*Danio rerio* (zebrafish) are a powerful model organism to study hematopoiesis (reviewed in 1-3). In recent decades, this model has permitted numerous studies of blood disorders. Zebrafish develop *ex utero* and are extremely fecund. Their ease (and low cost) of maintenance and husbandry, coupled with their genomic conservation with higher vertebrates allows inexpensive but thorough genetic assessments to be performed quickly and easily. Importantly, zebrafish are transparent during early embryonic stages which allow for *in vivo* visualization of development in real time. This is further enhanced by the plethora of blood-specific promoters that drive fluorescent protein expression and allow for live visualization of various blood cell types. Additionally, reverse genetic engineering methods such as clustered regularly intercepted short palindromic repeats RNA-guided Cas9 nucleases (CRISPR/Cas9) and transcription activator-like effector nucleases (TALENs) can be easily utilized for genetic manipulation, while forward genetic screens utilizing mutagenic chemicals and retroviruses have already elucidated a multitude of previously unknown pathways important for blood formation (4–15). Finally, zebrafish are an excellent model for large scale drug screens to treat and prevent hematopoietic disease (16–20). In fact, many drugs have been discovered through screens of zebrafish that are now in clinical trials for human use.

Previous work in our laboratory identified highly expressed transcripts in hematopoietic supportive cell lines (21–22). *rapzunel5* (*rpz5*) was among those genes highly expressed in zebrafish embryonic stormal trunk cells (ZEST; 23), kidney stromal cells (ZKS; 24), and caudal hematopoietic embryonic stromal tissue (CHEST; 21); all essential sites for regular hematopoiesis. Importantly, all of these cell lines support hematopoiesis *in vitro*, so they are excellent model systems to study signals necessary for HSPC support, proliferation, and differentiation.

*rapunzel* was first identified in an ENU mutagenesis screen, with the mutant having a skeletal overgrowth phenotype (25). The homozygous mutant embryos were also lethal at 4-5 dpf, which may have been from body axis deformities, pericardial edema, or poorly formed fin folds and jaws (25). Importantly, they also had defects in their vasculature and hematopoiesis, although these defects were not described in any detail (25). In the ENU mutated area of the genome were three paralogous genes, which were named *rpz*, *rpz2*, and *rpz3*. Two other paralogues (*rpz4* and *rpz5*) were found in other downstream areas of the genome after sequence comparisons were performed. Although not detected in mammals, *rpz5* plays an important role in the interferon response of teleosts (26). In this study, we describe a novel function of *rpz5* in the development of the immune system during development. Our data indicate that *rpz5* is essential for normal erythroid, myeloid, and thrombocyte formation, as well as HSPCs. This is specific to the erythro, myeloid, and thrombocytic lineages, however, as lymphoid cells were unaffected by the loss of *rpz5*.

## Materials and Methods

### Zebrafish husbandry and care

Zebrafish were mated, staged, and raised as described (27) and maintained in accordance with California State University, Chico Institutional Animal Care and Use Committee (IACUC) guidelines. All procedures were approved by the IACUC before being performed. Personnel were trained in animal care by taking the online Citi Program training course entitled “Working With Zebrafish (*Danio rerio*) in Research Settings” (https://www.citiprogram.org). Wildtype (AB) and the transgenic zebrafish lines *mpx*:EGFP (28), *itga2b*:EGFP (commonly referred to as *cd41:EGFP)* (29)*, lcr:EGFP* (30), and *lck*:EGFP (31) were used for these studies. Zebrafish were housed in a 700L recirculating zebrafish aquarium system (Aquatic Enterprises, Seattle, WA) regulated by a Profilux 3 Outdoor module that regulated salinity, pH, and temperature (GHL International, Kaiserslautern, Germany) 24-hours-a day. The facility was illuminated on a 14-hour light/ 10-hour dark cycle. Zebrafish were fed once a day with hatched brine shrimp (Brine Shrimp Direct, Ogden, UT) and Size 4 Zebrafish Research Diet (daniolab, Boston, MA).

### Generation of *rpz5* mRNA

*rpz5* transcript was amplified from zebrafish kidney cDNA using the following *rpz5* primers: FWD 5’-ATGAATCGGGTGGAAGAATGGG-3’ and REV 5’-CTATTCAGAGTGAATGCACAGG-3’. *rpz5* transcript was cloned into a TOPO-TA vector (Invitrogen, Carlsbad CA) and validated by Sanger sequencing. *rpz5* was then subcloned into pCS2+ and linearized with Not1. *rpz5* mRNA was generated using a mMessage SP6 kit (Ambion, Austin, TX).

### *rpz5* sequence read counts

Sequenced RNA libraries generated from zebrafish stromal cell lines were processed and analyzed as previously described (21, 23).

### Morpholino and *rpz5* mRNA injections

*rpz5* antisense morpholino (MO) was designed against the start codon to prevent translation of *rpz5* mRNA (Gene Tools, Philomath, OR). The MO sequence is as follows: 5’-AAACCCATTCTTCCACCCGATTCAT-3’. A control MO was also used with the sequence of 5’-CCTCTTACCTCAGTTACAATTTATA-3’. For microinjection into embryos, a mix of 3 μL of 8.26 mg/mL *rpz5* MO was mixed with 1 μL of phenol red for a final concentration of 6.2 ng/nL *rpz5* MO. 1 μL of this mix was loaded into a needle made with a PM102 micropipette puller (MicroData Instrument, Plainfield, NJ). Single-cell stage embryos were collected, placed onto a 1% agarose microinjection chamber plate with troughs, and injected with 1 nL (6.2 ng) of *rpz5* MO with a PM 1000 Cell Microinjector (MicroData Instrument, Plainfield, NJ). For rescue injections, rescued embryos received 6.2 ng of *rpz5* MO and 115 ng *rpz5* mRNA.

### Microscopic visualization of the hematopoietic system

Transgenic zebrafish were visualized under a Leica M165C (Leica, Wetzlar, Germany) fluorescent dissecting microscope at time points correlated with the emergence of specific hematopoietic cell lineages. Erythrocytes were visualized at 48 hpf with *lcr*:EGFP transgenic animals. Myeloid cells were visualized under the microscope at 48 hpf with *mpx*:GFP transgenic animals. Thrombocytes were visualized with *cd41*:EGFP transgenic fish at 72 hpf. T cells were visualized at 120 hpf with *lck*:GFP transgenic fish. Zebrafish and their fluorescently labeled cells were imaged using a Leica FireCam camera (Leica, Wetzlar, Germany), scored, and enumerated. For myeloid quantitation, images were labeled by a reference number and the numbers of *mpx*:GFP+ cells were counted in each animal by several undergraduate students to ensure no bias in results. Results were validated with a machine learning program that was trained to count fluorescent cells (32). For T cell quantitation, images of *lck*:GFP embryos were analyzed with ImageJ (https://imagej.nih.gov/ij/ ; 33) to determine the pixel density of each thymus relative to background.

### Quantitation of HSPCs in developing zebrafish embryos

HSPC isolation and culture was performed as previously described (34–36). Samples had carp serum, GCSF, and EPO added to stimulate myeloid and erythroid differentiation (35, 37). They were incubated at 32°C and 5% CO2 for 7–10 days and imaged with an Olympus IX53 inverted microscope (Olympus, Center Valley, PA) at 20x to enumerate colony forming units (CFUs).

### rpz5 PCR

mRNA was extracted from embryos at 24 and 48 hpf using a Qiagen RNAeasy kit (Qiagen, Hilden, Germany). cDNA was then generated with the iScript cDNA synthesis kit (Biorad, Hercules, CA), and PCR was performed with JumpStart Taq DNA polymerase (MilliporeSigma, Burlington, MA). *ef1a* (38) was used as a reference gene.

### Statistics

To discern statistical differences, data were analyzed using an unpaired two-tailed Student’s T test with Microsoft Excel.

## Results

To identify novel transcripts likely involved in hematopoiesis, we examined the transcriptome of three hematopoietic-supportive zebrafish cell lines (ZKS, ZEST, and CHEST cells) and compared the transcripts to a non-supportive cell line (ZF4 cells) (see 21, 23). *rpz5* levels were 8.9x higher in CHEST cells, 20.2x higher in ZEST cells, and 30.7x higher in ZKS cells (**Table 1**) when compared to ZF4 cells. This elevated amount of *rpz5* in these cell lines indicated that the transcript was likely involved in hematopoiesis. To verify that *rpz5* was expressed in developing zebrafish embryos, PCR was performed at 24 and 48 hpf. *rpz5* transcript was detectable at 48 hpf, indicating that we could modulate its activity and see the downstream effect on blood cell formation (**Fig 1A**). To verify if we could reduce *rpz5* levels with a specific MO, we generated a MO that targeted the gene’s translational start site. Injection of no MO and control MO had no effect on the levels of *rpz5*, while the specific *rpz5* MO entirely prevented *rpz5* expression, likely due to mis-splicing or nonsense-mediated decay (**Fig 1B**).

**Fig 1:**
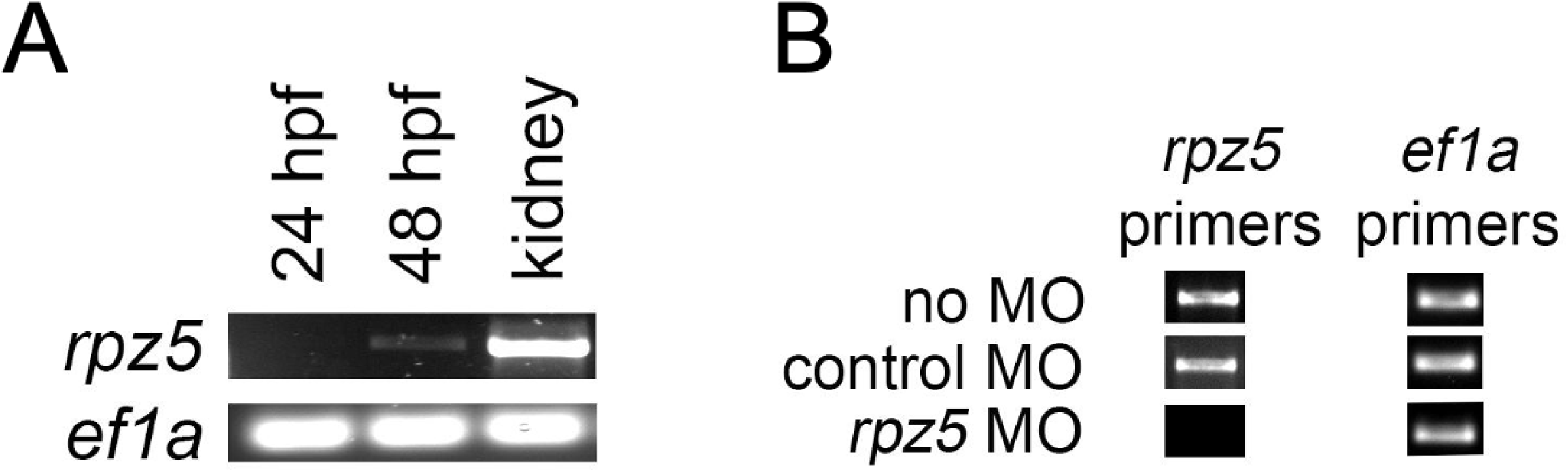
*rpz5* is expressed in the developing zebrafish embryo and reduced by *rpz5* MO. **(A)** 10 whole zebrafish embryos were digested, RNA was extracted, cDNA generated, and subjected to RT-PCR with primers for *rpz5* (top) and *ef1a* (bottom) at 24 hpf and 48 hpf. Kidney is shown as positive control. **(B)** Embryos were injected at the one-cell-stage with PBS (no MO), control MO, or *rpz5* MO. 10 whole zebrafish embryos were digested, RNA was extracted, cDNA generated, and subjected to RT-PCR with primers for *rpz5* (left) and *ef1a* (right) at 48 hpf.

**Table 1:** *rpz5* is highly expressed in hematopoietic-supportive stromal cell lines. Each stromal cell line is derived from sites of hematopoietic emergence, expansion, and maintenance in adult and embryonic zebrafish; adult zebrafish kidney stroma (ZKS, derived from the adult kidney), zebrafish embryonic stromal trunk (ZEST, derived from the embryonic trunk), and caudal hematopoietic embryonic stromal tissue (CHEST, derived from the CHT). Each stromal cell line’s transcriptome was analyzed with RNA-seq and compared to a non-hematopoietic supportive stromal cell line (ZF4); transcript levels for each cell line are presented as reads per kilobase of transcript per million reads mapped (RPKM).

| **Stromal cell lines** | **Transcript levels (RPKM)** |
| --- | --- |
| Zebrafish kidney stromal (ZKS) cells | 239.96 |
| Zebrafish embryonic stromal trunk (ZEST) cells | 157.82 |
| Caudal hematopoietic embryonic stromal trunk (CHEST) cells | 69.99 |
| Zebrafish fibroblast 4 (ZF4) cells (non-supportive) | 7.81 |

To examine the effect of *rpz5* on morphology, *rpz5* MO was injected at the one-cell-stage and embryos were allowed to develop until 48 hpf. Most embryos were morphologically normal (**Fig 2A**), but a few had pericardial effusion (**Fig 2B**).

**Fig 2.**
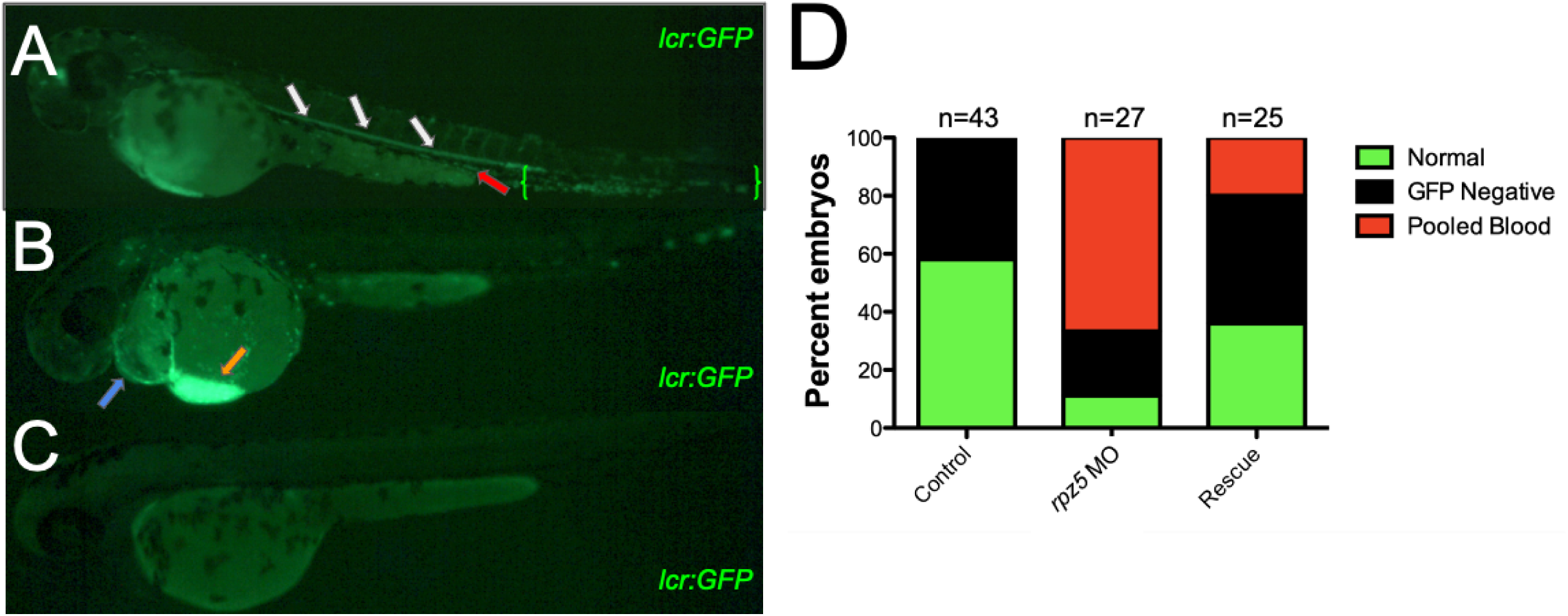
*rpz5* knockdown reduces erythrocytes in circulation. *lcr:GFP* zebrafish embryos were injected at the single-cell-stage with 6.2 ng *rpz5* MO (rpz5 MO), or 6.2 ng *rpz5* MO and 115 ng of *rpz5* mRNA (Rescue); uninjected embryos served as controls (Control). 24 hpf zebrafish visualized at 50x that have normal **(A)**, pooled **(B)**, and negative GFP **(C)** blood. White arrows indicate the dorsal aorta and red arrows indicate the cardinal vein. Green brackets show the caudal hematopoietic tissue, the orange arrow indicates blood pooling of blood in the Ducts of Cuvier, and blue arrow denotes pericardial effusion. **(D)** Quantitation of zebrafish shown in (**A-C**); pooled blood (red), negative GFP (black), and normal (green) phenotypes. Numbers (n) of embryos examined are shown over the bars.

To observe if this loss of *rpz5* had a negative effect on blood formation, we first focused on red blood cells, one of the first blood cells produced. *lcr*:GFP fish, which have the locus control region of alpha globin that drives GFP fluorescence, allow all erythrocytes to be visualized. *rpz5* morphant zebrafish were categorized into circulation phenotypes of “normal” (**Fig 2A**), “pooled” (**Fig 2B**), or “negative” (**Fig 2C**). While “negative” fish had no labeled erythroid cells, “pooled” circulation refers to a buildup of erythrocytes especially in the anterior region of the embryo near the heart of the *lcr*:GFP zebrafish at 48 hpf. In this phenotype, the heart is still pumping, and liquid can be seen in the vessels of the animal. Approximately 60% of *rpz5* knockdown fish exhibited significant pooling of blood compared to none in the control group (**Fig 2D**). This abnormal circulation phenotype was reduced to about 20% in the rescue zebrafish. At the same time, fish with “normal” blood were reduced from approximately 60% to 10% when *rpz5* MO was added (**Fig 2D**). *rpz5* mRNA again rescued that phenotype, increasing numbers to almost 40%. Given these data, we suggest *rpz5* knockdown induced pooling of erythrocytes, reducing the number of cells in circulation.

Besides erythrocytes, macrophages and neutrophils are also some of the earliest blood cells produced during development. To determine the role of *rpz5* on these cell types, myeloid-specific transgenic animals (*mpx*:GFP) were injected with *rpz5* MO. At 48hpf, morphant and control zebrafish were visualized and imaged (**Fig 3A**). The total fluorescent myeloid cell count was determined by multiple undergraduate students and confirmed via a machine learning neutrophil counting model (32). *rpz5* morphant fish displayed a significant reduction in *mpx*:GFP cells compared to the control group (**Fig 3B**). Rescue experiments with *rpz5* mRNA recovered the numbers of *mpx*:GFP^+^ myeloid cells, confirming that *rpz5* plays an important role in myelopoiesis.

**Fig 3.**
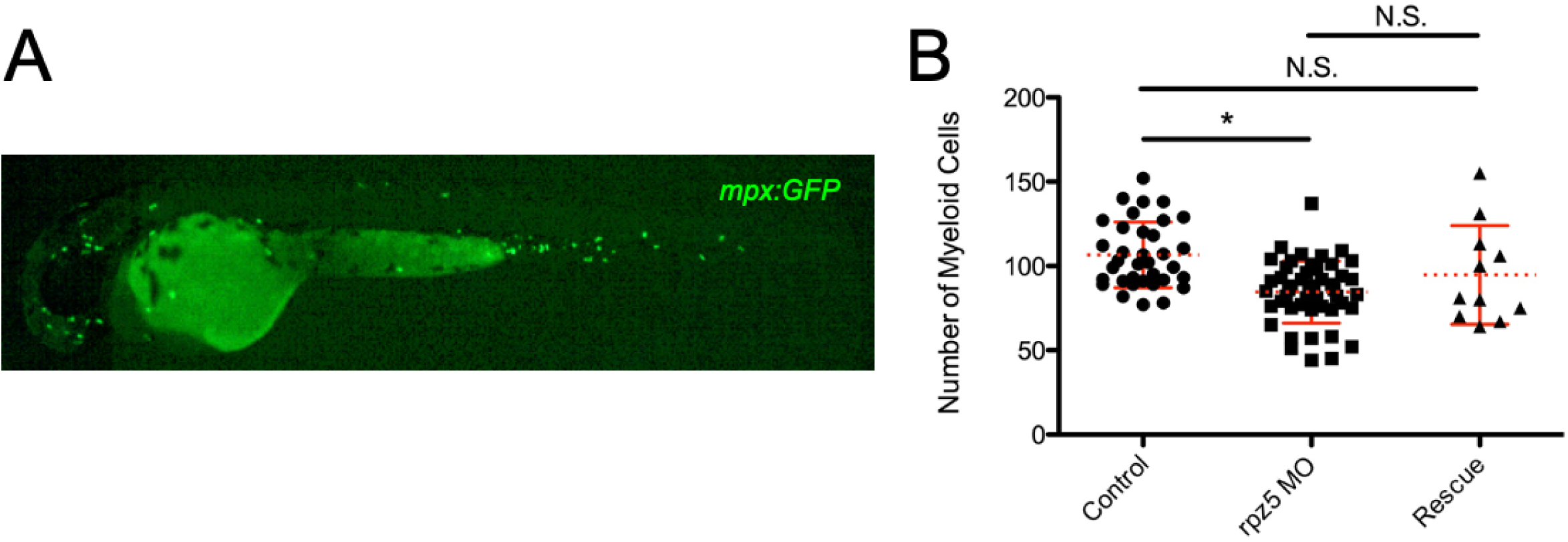
*rpz5* knockdown reduces myeloid cells. *mpx:GFP* zebrafish embryos were injected at the single-cell-stage with 6.2 ng *rpz5* MO (rpz MO, squares), or 6.2 ng *rpz5* MO and 115 ng of *rpz5* mRNA (Rescue, triangles); uninjected embryos served as controls (Control, circles). 48 hpf zebrafish were visualized and imaged at 50x **(A)** Representative *mpx:*GFP embryo at 48 hpf. Every green spot denotes an individual myeloid cell. **(B)** Fluorescent myeloid cells were manually quantitated for each condition. Dashed line denotes median, and bars represent SD. * denotes p < 0.0001, and N.S. denotes no significance.

Zebrafish erythrocytes and thrombocytes share a common progenitor cell called the thrombocyte erythroid progenitor (TEP). Due to the fact that circulating erythrocytes were reduced, we suspected that thrombocyte production might also be negatively impacted. To perform these studies, we utilized *cd41*:GFP transgenic animals. After injection, fish were allowed to develop for 72 hours, which is how long thrombocytes take to develop. The fish were then scored as “normal” (**Fig 4A**), “reduced” (**Fig 4B**), or “negative” (**Fig 4C**), depending on how many thrombocytes were visible. The “reduced” phenotype included transgenic fish with less thrombocytes in the caudal hematopoietic region; these fish also had slowed circulation. In the *rpz*5 MO KO group, 40% of fish had reduced numbers of thrombocytes, and only 25% normal amounts when compared to control (**Fig 4D**). Rescue with mRNA rescued this, with only 10% fish having reduced thrombocytes and normal fish increasing to 40%. Overall, *rpz5* is also required for the normal formation of thrombocytes in the developing zebrafish embryo.

**Fig 4.**
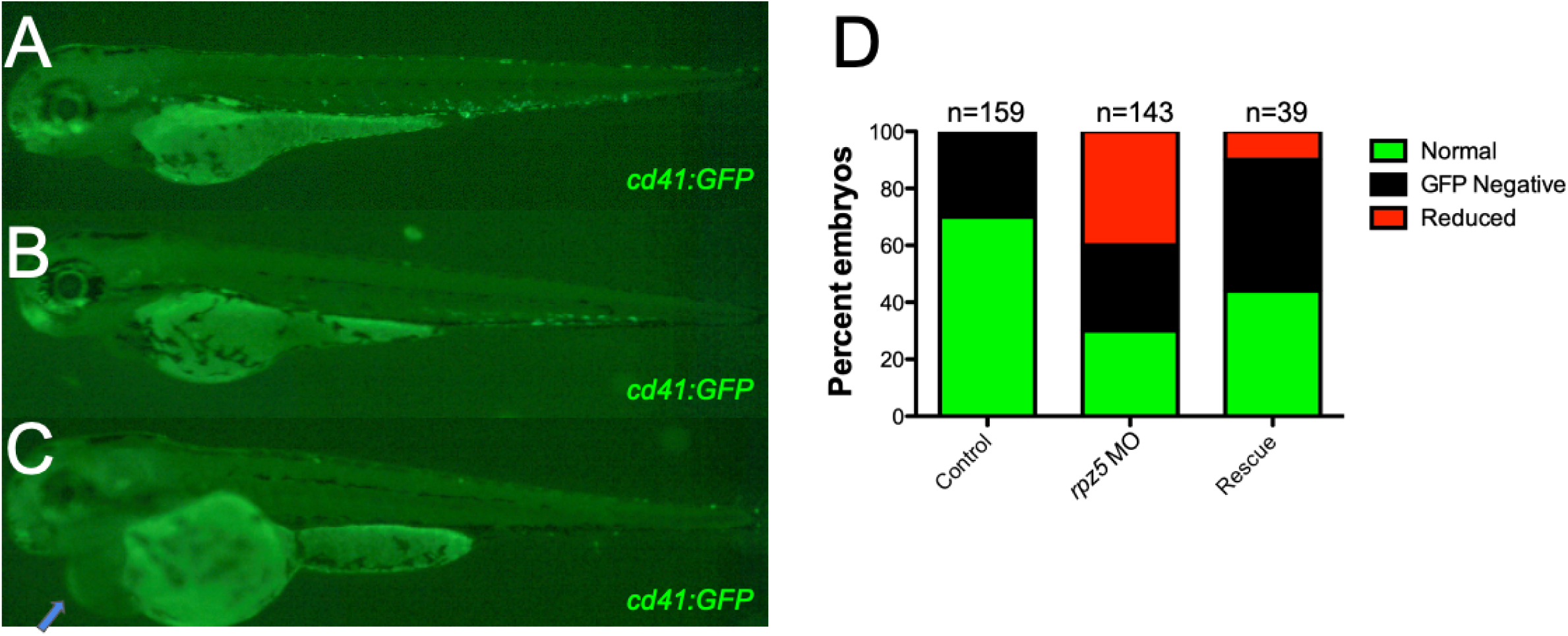
*rpz5* knockdown reduces thrombocytes. *cd41:GFP* zebrafish embryos were injected at the single-cell-stage with 6.2 ng *rpz5* MO (rpz5 MO), or 6.2 ng *rpz5* MO and 115 ng of *rpz5* mRNA (Rescue); uninjected embryos served as controls (Control). 72 hpf zebrafish visualized at 50x that have normal **(A)**, pooled **(B)**, and negative GFP **(C)** blood. Blue arrow denotes pericardial effusion. **(D)** Quantitation of zebrafish shown in (**A-C**); reduced thrombocytes (red), negative GFP (black), and normal (green) phenotypes. Numbers (n) of embryos examined is shown over the bars.

Lymphoid cells are another type of hematopoietic cells that are essential for the adaptive immune system. While B cell transgenic animals exist, their fluorescence is difficult to discern until later in development (39). To examine T cells, we utilized a transgenic animal where the *lck* promoter is driving GFP fluorescence. To perform these experiments, fish were injected with *rpz5* MO or control MO and allowed to develop for 5 days. At 5dpf, animals’ thymi were imaged at the same magnification and exposure settings (**Fig S1A-C**). Afterwards, pixel density of the thymus was quantified with ImageJ (**Fig S1D**). Control and morphant animals had no difference in thymic fluorescence, indicating that they had similar numbers of T cells and *rpz5* loss had no negative effects on T cell development.

All hematopoietic cells derive from a hematopoietic stem cell (HSC) that differentiates into more lineage-restricted hematopoietic stem and progenitor cells (HSPCs). While there are many ways to measure HSPCs in the developing embryo, we utilized a methylcellulose assay to quantitate these progenitors. To determine the impact of *rpz5* on HSPCs, pools of control and *rpz5* MO injected animals were collected at 48 hours, enzymatically digested, and plated in methylcellulose with supportive cytokines (**Fig 5A**). The samples were incubated for 7 days, after which the resultant colonies were visualized and counted. Colony forming units (CFUs) for *rpz5* morphant zebrafish were reduced on average by about 40% compared to the control group (**Fig 5B**). This reduction was completely rescued by mRNA injection (**Fig 5B**). These data suggest that *rpz5* is involved in the normal formation of HSPCs, which likely affects the downstream levels of erythroid, myeloid, and thrombocytic cells in the developing zebrafish.

**Fig 5.**
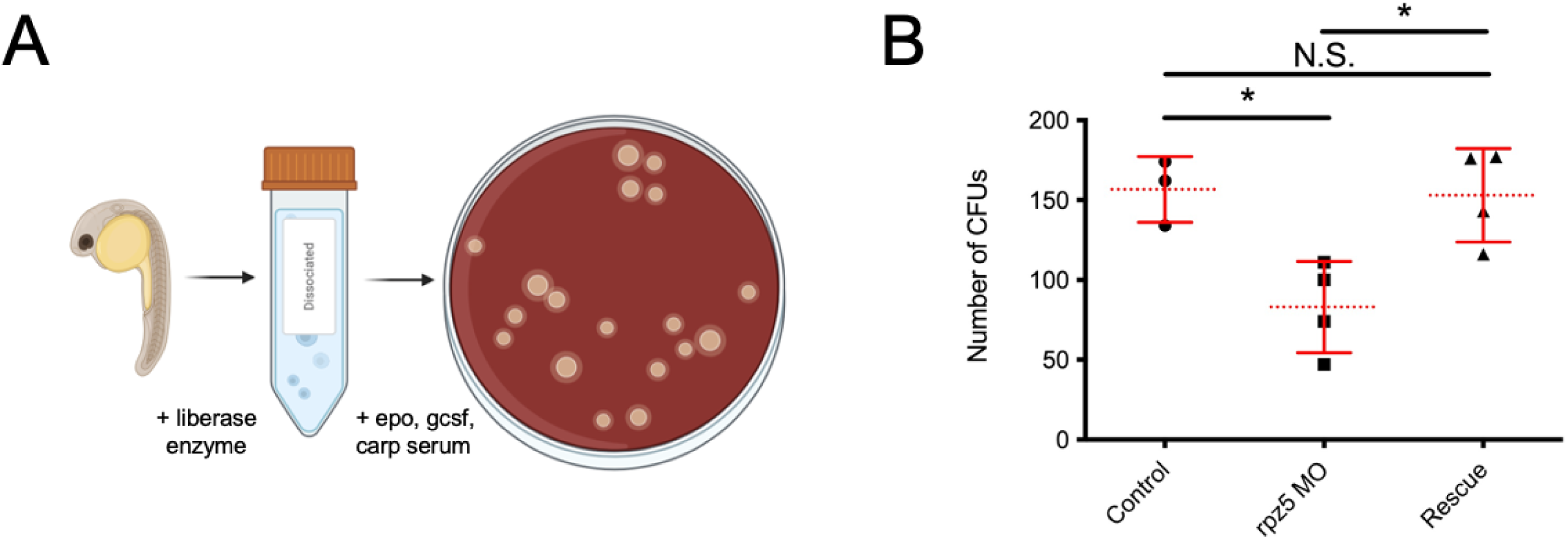
*rpz5* knockdown reduces HPSCs. Zebrafish embryos were injected at the single-cell-stage with 6.2 ng *rpz5* MO (rpz MO, squares), or 6.2 ng *rpz5* MO and 115 ng of *rpz5* mRNA (Rescue, triangles); uninjected embryos served as controls (Control, circles). 48 hpf zebrafish were digested and subjected to a methylcellulose assay **(A)**. Images created with biorender.com **(B)** Colony forming units (CFUs) from each condition are shown. Each point denotes 12 randomly selected animals that were digested and plated. Dashed line denotes median, and bars represent SD. * denotes p < 0.02, and N.S. denotes no significance.

## Discussion

Hematopoiesis is an extremely complex process necessary for the survival of an organism. This process begins in the early embryo, with primitive erythroid and myeloid cells arising around 24hpf. Then, definitive HSPCs and HSCs arise, which will give rise to all hematopoietic cells, including lymphoid lineages, for the remainder of the animal’s life (reviewed in 40-42). While research on hematopoiesis has occurred for decades, many of the molecular pathways responsible for the differentiation of these HSCs and HSPCs have yet to be elucidated. To address that issue, we isolated hematopoietic-supportive cell lines and interrogated their transcriptome, with the hopes that they were producing and secreting hematopoietic-supportive proteins. One of the proteins that we detected from these supportive cell lines was *rpz5*.

RPZ5 belongs to the *rpz* family, composed of *rpz*, *rpz2*, *rpz3*, and *rpz4* in zebrafish. While *rpz5* has not been detected in mammals, it is well conserved in teleosts. The zebrafish mutant *rapunzel* (a missense mutation of *rpz*) results in skeletal overgrowth (25) even though the proteins’ actual molecular function is unclear. To the best of our knowledge the other family members have not been examined, excluding *rpz5*, which is a negative regulator of RIG-I like receptor (RLR)-mediated interferon (IFN) production (26). In other words, Lu et al found that *rpz5* played an important role in the fish’s response to viral infection. There are two transcript variants of *rpz5*, and while we likely reduced both versions with our 5’ MO designed to the transcriptional start site, we saw effective rescue with the long transcript (rpz5-201 in www.ensembl.org), which codes for a 393aa protein that is ∼45kDa. Our findings, that *rpz5* is involved in the early development of the immune system, fits with Lu et al.’s previous work. Proinflammatory signaling is essential in the early formation of the hematopoietic system, especially during the emergence of HSCs during development (43–45) and IFN plays an important role in the proliferation of HSCs, preserving self-renewal and multilineage differentiation capability (46 and reviewed in 47-48). The reduction of IFN signaling due to *rpz5* knockdown may directly explain why less HSCs and HSPCs are seen in morphants. Additionally, while *rpz* mutants had skeletal malformations (25), our MO resulted in no overt phenotypical skeletal issues. It appears that *rpz5*, either directly or through IFN signaling, has a novel role in the development of the hematopoietic system.

In this study we knocked down *rpz5* with a MO. MOs are excellent methods to reduce the expression of a protein, especially during development. When we knocked down *rpz5*, we saw abnormal erythroid, myeloid, and thrombocytic differentiation. Rescue with *rpz5* mRNA effectively rescued these defects, indicating that phenotypes were likely directly caused by *rpz5* reduction. While CRISPR-Cas9 mutants could be produced, there is a strong chance that one of the other four family members, if they have any functional redundancy, could be overexpressed and mask the knockout’s effect (25, 49). The effects are clearly not off-target, as T cells were not reduced. Overall, these findings further validate the use of MOs to discern important biological processes, especially during development.

As mentioned above, the fact that *rpz5* reduction has no effect on lymphoid cells is interesting. HSCs are the keystone cells of the hematopoietic system, which further differentiate into common lymphoid progenitors (CLPs; 50), and common myeloid progenitors (CMPs; 51). Further downstream of CMPs are granulocyte monocyte progenitors (GMPs; 51) and thrombocyte erythrocyte progenitors (TEPs; 52). Using a methylcellulose assay adapted for zebrafish, we saw that HSPC numbers were reduced. As we only added granulocyte colony stimulating factor (gcsf), erythropoietin (epo), and carp serum to the mix, we can only really assess CMPs and GMPs, as those cytokines are essential for those HSPCs to differentiate (35–36). Thrombopoietin could be added into the media, but it would likely have no effect, as thrombocytes were reduced in *rpz5* morphants and TEPs are downstream of CMPs and GMPs. While lymphoid cells are notoriously hard to examine *in vitro*, these studies indicate that *rpz5* likely affects downstream HSPCs like the CMP and GMP, but not necessarily HSCs; if HSCs were negatively affected, we would expect less T cells present.

Overall, these studies describe a novel role for *rpz5* in hematopoietic development, indicating that it is important for normal erythro-, thrombo-, and myelo-poiesis while appearing seemingly dispensable for T cell differentiation. While this gene is not present in mammals, it is well conserved in teleosts. Further study of this gene is needed to understand it’s role in hematopoiesis, but it is likely functioning through down-regulating IFN signaling, which affects HSPC formation and differentiation.

---

Fig S1. *rpz5* knockdown has no effect on T cells.

*lck:GFP* zebrafish embryos were injected at the single-cell-stage with 6.2 ng *rpz5* MO (rpz5 MO, squares); uninjected embryos served as controls (control, circles). 120 hpf zebrafish visualized at 50x that have normal **(A)**, low **(B)**, and negative GFP **(C)** thymi (red dotted circle), where T cells are formed. * denotes background fluorescence of the yolk ball. **(D)** Quantitation of thymic pixel density shown in (**A-C**) with Image J. Dashed line denotes median, and bars represent SD. N.S. denotes no significance.

